# Detecting and harmonizing scanner differences in the ABCD study - annual release 1.0

**DOI:** 10.1101/309260

**Authors:** Dylan M. Nielson, Francisco Pereira, Charles Y. Zheng, Nino Migineishvili, John A. Lee, Adam G. Thomas, Peter A. Bandettini

## Abstract

In order to obtain the sample sizes needed for robustly reproducible effects, it is often necessary to acquire data at multiple sites using different MRI scanners. This poses a challenge for investigators to account for the variance due to scanner, as balanced sampling is often not an option. Similarly, longitudinal studies must deal with known and unknown changes to scanner hardware and software over time. In this manuscript, we have explored scanner-related differences in the dataset recently released by the Adolescent Brain Cognitive Development (ABCD) project, a multi-site, longitudinal study of children age 9-10. We demonstrate that scanner manufacturer, model, as well as the individual scanner itself, are detectable in the resting and task-based fMRI results of the ABCD dataset. We further demonstrate that these differences can be harmonized using an empirical Bayes approach known as ComBat. We argue that accounting for scanner variance, including even minor differences in scanner hardware or software, is crucial for any analysis.

## Introduction

The Adolescent Brain Cognitive Development (ABCD) Study is a multisite, longitudinal study designed to recruit more than 10,000 children age 9-10 and follow them over 10 years into early adulthood (Casey et al., 2018). The project has recently released its first installment of 4,094 participants collected at 21 different sites across the United States (Jernigan et al., 2018). The ABCD project represents the latest advance in large, collaborative neuroimaging initiatives. It joins other high impact projects such as the Alzheimer’s Disease Neuroimaging Initiative (ADNI), the Human Connectome Project (HCP), and the UK Biobank Imaging initiative (Van Essen et al., 2013; Jack et al., 2008; Miller et al., 2016). The ABCD project has set a new standard for rapid, coordinated data sharing by making both the raw images and preprocessed data available well before the primary investigators had analyzed or published on them.

Collecting a data set of this caliber across multiple sites is a demanding undertaking (Toga and Dinov, 2015). Among the challenges is the harmonization of scanner sequences. The 21 ABCD sites include scanners from three different vendors and several different models; there are unavoidable differences in the data they produce. Scanner manufacturers typically optimize scanners for a medical setting, i.e. to be viewed and interpreted by a trained physician. Thus it falls to the site physicists and investigators to calibrate scanners so as to produce standardized image data suitable for scientific inquiry. This is further complicated by closed source, proprietary reconstruction software preventing an understanding of how the image data from a given scanner is derived.

Several studies have explored the effects of scanner variance on multi-site studies and have reached different conclusions. Some authors have concluded that carefully harmonized scanner sequences have a minor effect on multi-site analyses that are outweighed by the benefits (Noble et al., 2017). Others have found scanner effects to be a major confound that sometimes shows interactions with variables of interest such as age (Focke et al., 2011; Auzias et al., 2016). The genetics community has been faced with similar challenges in the analysis of microarray data, for which different batches may give different results due to a variety of factors within and between labs. ComBat (COMbining BATches) is a popular empirical Bayes approach to correcting these batch effects (Johnson et al., 2007), which works by obtaining an initial estimate of batch interaction parameters from a linear model, then shrinking those interaction parameters towards a grouped mean, with a level of shrinkage that is calibrated based on either parametric or nonparametric estimates of the distribution of batch effects. This approach has also been shown to be an effective tool for correcting cross-site effects in human neuroimaging for DTI data (Fortin et al., 2017) and cortical thickness measures (Fortin et al., 2018).

The first goal of this paper is to propose a quantitative test of whether there are systematic differences between scanners. The second goal is to evaluate an approach for eliminating or attenuating those differences in the functional MRI data in the recently released ABCD dataset (Jernigan et al., 2018).

Our test is based on the intuition that if systematic differences between scanners exist, it should be possible to predict the scanner that each subject’s imaging data was collected on. We do this using a straightforward logistic regression classifier, trained on a variety of quantitative measures derived from imaging data. Conversely, such a classifier should lose the ability to make this prediction if the differences were successfully eliminated. To do this we use ComBat (Johnson et al., 2007; Fortin et al., 2018) to identify and remove mean and variance differences between corresponding variables in comparable samples. In our case, samples contain participants scanned on a given scanner, and the variables collected for each participant are functional connectivity values or task-based average beta weights from a set of ROIs.

## Methods

### Data

For our analyses we used the pre-computed resting state and task-based metrics contained in ABCD’s annual release (Jernigan et al., 2018). Generation of the resting state and task-based metrics are described in the release notes (Jernigan et al., 2018). ABCD’s task-based and resting state fMRI (rs-fMRI) data includes information from 4,094 participants (2,140 male) with a mean age of 120.1 months (*SD* = 7.35 months). Each subject was scanned on one of 28 different different scanners, identified by their hashed serial number. Information on where each of these 28 scanners were sited is not provided. Based on ABCD’s extensive quality control metrics there are 2,852 participants with acceptable rs-fMRI data. ABCD participants performed three fMRI tasks: Monetary Incentive Delay (MID), Stop Signal Task (SST), and nBack. 1,997 participants have data that passes ABCD’s quality control metrics on two runs of each of these tasks. The rs-fMRI metrics consist of connectivity within and between 12 cortical resting state communities plus parcels not assigned to any community as well as the connectivity between these networks and 19 subcortical ROIs (Gordon et al., 2016). The 12 networks are: Auditory, Cingulo-Opercular, Cingulo-Parietal, Default, Dorsal Attention, Fronto-Parietal, Retrosplenial-Temporal, Salience, Somatomotor Hand, Somatomotor Mouth, Ventral Attention, and Visual. “None” refers to ROIs not assigned to any of the aforementioned communities. ABCD has made available beta weights and T-statistic values for various conditions and contrasts, averaged within each of the 333 ROIs defined in Gordon et al. for each subject. In our analyses of the task-based data we use the averaged beta weights from each ROI. Information about participant handedness comes from the ABCD Youth Edinburgh Handedness Inventory Short Form (EHIS; Oldfield, 1971). We limited our analysis of rs-FMRI to the 13 scanner IDs that had scans passing quality control for at least 100 participants, leaving 2,816 participants (1,471 Male). Our analysis of the task-based data is limited to the 7 scanner IDs with at least 100 participants with 2 runs for each task passing ABCD’s quality control metrics, leaving 1,069 subjects.

### Cross-validation and sample balancing

We fit and tested scanner classification models on both rs-FMRI and task-based data, using three-fold cross-validation in each case. In order to ensure that classification was not driven by imbalances in the distribution of sex and handedness across scanners, we used a balancing procedure. For each combination of subject characteristics, the same number of subjects was used across all scanners; if more were available for a particular scanner, the correct number was drawn at random. We produced 25 different balanced datasets, and ran the whole cross-validation procedure for each. In the rs-fMRI data, these balanced datasets had 89 subjects from each of 13 scanners for a total of 1,157 subjects per draw. The task-based data had 87 subjects from each of 7 scanners for a total of 609 subjects per draw. The results reported are the average of results obtained using each of the twenty five datasets.

In addition to balancing sex and handedness, we tried two different approaches to control for differences in age distributions between scanners. First, we controlled for age effects by regressing age out of each metric individually. Regressing out age effects may not eliminate them completely, so we also extended the balancing procedure to age in three month bins, in addition to sex and handedness as described above. In the rs-fMRI data each sex, handedness, and age balanced draw had 27 subjects from each of 13 scanners for a total of 351 subjects per draw. It was not practical to draw age balanced datasets for the task-based data given the relatively low number of subjects per scanner, so for these analyses we only regressed the effect of age out of each metric.

### Predicting Scanner

We predicted scanner separately for the rs-fMRI data and each run of each contrast or condition from the task-based data. We performed all of these classifications with multinomial logistic regressions with an L2 penalty, a regularization strength of 1 and the SAGA (Defazio et al., 2014) solver with 10,000 maximum iterations. The L2 penalty and regularization strength are defaults in Scikit-Learn (Scikit-learn, 2011). We fit a multinomial classifier instead of one-versus-rest to simplify interpretation of the classifiers’ weights. While other choices were certainly possible, our primary goal is to test the hypothesis that scanner differences in rs-fMRI and task data exist, rather than build the best possible scanner classifier.

Metrics were z-scored based on each training split prior to fitting the logistic regression. When we regressed the effect of age out of each metric, the linear model of age predicting that metric was fit on the training split and the same coefficients were used to correct the test set. Additionally, we investigated the ability of ComBat (Johnson et al., 2007) to eliminate the variance facilitating classification of scanner. We estimated ComBat parameters separately for each training split and test split. We determined chance classifier performance with 1,000 sex, handedness and age balanced permutations for the rs-fMRI data and 1,000 similarly balanced permutations for the task-based data. Age was balanced in three month bins. These permutations were also used to define null distributions for cells of the confusion matrices examined for rs-fMRI data. We assessed differences in true-positive rate (TPR) for scanners from each manufacturer with pairwise t-tests of the mean TPR across manufacturer from all draws. Significance controlling for multiple comparisons was determined with a null distribution constructed from the most extreme pairwise t-value from each permutation. The ability of ComBat to reduce performance on each scanner was calculated as the reduction in TPR for each scanner and compared to null distributions of reductions and increases in TPR from the permutations. Multiple comparisons correction was achieved by taking the greatest reduction and greatest increase from each permutation to build the positive and negative tails of the null distribution, respectively.

### Percent variance explained

We examined the percent variance in each rs-fMRI and task-based metric explained by scanner manufacturer, model, and serial number. The percent variance quantifies the impact of scanner differences on the rs-fMRI and task-based data on a scale useful for determining the importance of these effects. This nuisance variance may, in the worst case, interfere with the detection of effects of interest, and should be accounted for in any modeling done with these variables. We calculated percent variance due to these factors as the increase in *R*^2^ when adding each factor to the model predicting each metric. For calculation of variance explained we used the entire sample of 4,094 participants for the rs-fMRI and 1,997 participants for the task-based data,with no balanced draws. We used the entire sample here so that we could get the fullest picture of the contributions of these factors to the percent variance explained.

We fit the following 5 models for each metric provided in the rs-fMRI and task-based data, using ordinary least squares:

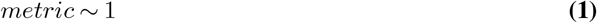

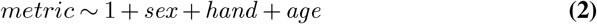

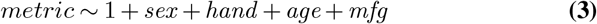

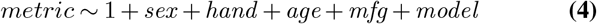

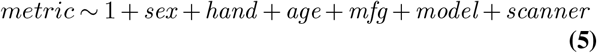

Thus we get the percent variance explained by sex, handedness, and age by subtracting the percent variance explained by equation 1 from the percent variance explained by equation 2, the percent variance explained by manufacturer by subtracting the percent variance explained by equation 2 from the percent variance explained by equation 3, etc. Computing percent variance explained in this way will first assign any variance shared between two factors to the factor that is considered in the simpler model, i.e. if some metric’s variance is shared between model and manufacturer, that variance will count towards the variance explained by manufacturer.

We also tested the ability of ComBat to reduce the percent variance explained by manufacturer, model, and scanner ID. For these tests, ComBat parameters were fit based on the entire sample with no training-test split. We determined the null distributions for the percent variance explained with the same permutations used for predicting scanner. As described above, we balanced these permutations on sex, handedness, and age so that our null distribution takes into account the imbalance of these factors between scanners. This gives us a more accurate null distribution for testing the percent variance explained by manufacturer, model, and scanner, but it precludes assessing the significance of the percent variance explained by sex, handedness, and age.

### Spatial distribution of scanner effects

Scanner was strongly identifiable in all runs of nBack conditions, which provided an opportunity to look for consistent spatial patterns driving that identifiability. We ran a t-test of the null hypothesis that the mean value of each coefficient was 0 across all draws and cross-validation folds, for all of the nBack conditions. A coefficient was deemed consistent if the absolute value of the t-statistic for that coefficient was more extreme than a null distribution of the most extreme t-values from all coefficients in each permutation, giving a significance threshold of 0.002.

### Software

All code for the analysis along with instructions for reproducing the compute environment used can found at http://github.com/nih-fmrif/nielson_abcd_2018.git.

## Results

### Resting State Functional Connectivity

We classified collection scanner in the ABCD resting state data with a multinomial logistic regression with L2 normalization (Figure 1). Average accuracy over three-fold cross validation from 25 draws balanced for sex and handedness was 35.3% (*SD* = 2.1%, *p* < 0.002), well above chance performance as determined from 1,000 sex, handedness and age balanced permutations.

**Fig. 1.**
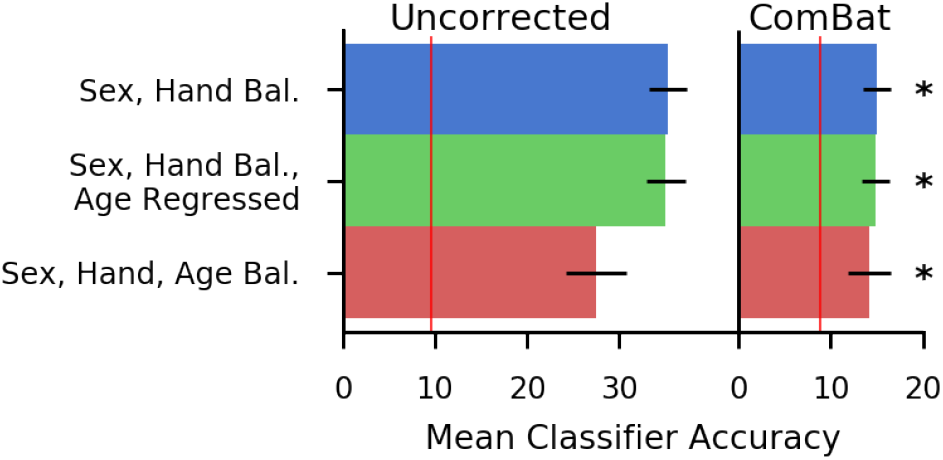
Classifier Performance Before and After ComBat for Resting State Functional Connectivity. Bars show mean classifier performance predicting collection scanner across three-fold cross-validation for raw and age regressed data and split half cross-validation for age balanced data from 25 gender and handedness balanced draws for resting state functional connectivity metrics. Error bars indicate standard deviation across draws and cross-validation folds. We ran 1,000 permutations balanced across gender, handedness and age in three month bins to determine chance classifier performance. The red lines indicate the multiple comparison corrected *p* < 0.002 threshold for chance performance. For the uncorrected bars the threshold is 9.6% for ComBat it is 8.9%. Bars with performance greater than this cut off are indicated with more saturated colors. Classifier performance on data with no corrections for age are shown in blue. Green bars show classifier performance in which the effect of age learned on the training split within each cross validation fold was regressed out of each ROI. Red bars show classifier performance using a restricted dataset in which we were able to draw samples balanced for gender, handedness and age in three month bins from each scanner. The uncorrected columns show results without running ComBat to correct for scanner effects. The ComBat columns show classifier performance after correcting for scanner effects with ComBat separately on the training and test data within each cross validation. Stars indicate that ComBat significantly reduced classifier performance with *p* < 0.002 threshold.

Performance was still above chance when controlling for age by either regressing the effect of age out of each factor (mean accuracy 35.3%, *SD* = 2.0%, *p* < 0.002), or when using data with sex, handedness, and age in three month bins balanced across scanners (mean accuracy 27.5%, *SD* = 3.2%, *p* < 0.002; Figure 1). We controlled for multiple comparisons by taking the maximum and minimum accuracies from each of the 3 permutations and using these as the upper and lower tails of the null distribution respectively. (Maris and Oostenveld, 2007). The p-value for each value is thus the lesser of its p-values when compared to the null distributions for each tail multiplied by two. The chance performance thresholds were 9.6% for uncorrected data and 8.9% for ComBat corrected data.

We evaluated ComBat as an approach to control scanner effects by applying it separately to the training and test data in each cross-validation split. Accuracy after ComBat correction was lower than without correction for sex and handedness balanced draws, age regressed sex and handedness balanced draws, and sex, handedness, and age balancing (*p* < 0.002 for all, calculated as above). However, classification after ComBat correction was still higher than chance for all three balancing approaches. Examining classifier performance more closely via the confusion matrices (Figure 2) shows that rs-fMRI data from scanners manufactured by Philips (mean TPR = 0.75, *SD* = 0.05) was more distinguishable that rs-fMRI data from GE (mean TPR: 0.37, *SD* = 0.04) and both were more distinguishable than data from Siemens scanners (mean TPR = 0.24, *SD* = 0.02) with *p* < 0.002 for all comparisons. In the sex, handedness, and age balanced rs-fMRI data, Philips (mean TPR = 0.61, *SD* = 0.01) was more distinguishable than GE (mean TPR = 0.29, *SD* = 0.07) or Siemens (mean TPR = 0.18, *SD* = 0.04) with *p* < 0.002. Aggregating these results to the model and manufacturer level we can see that the classifier is more likely to mis-classify data from one scanner as coming from another scanner with the same model, or from a scanner with the same manufacturer. ComBat reduced scanner identifiability with *p* < 0.002, but it did not entirely eliminate the tendency for scanners to be misclassified for other scanners with the same manufacturer or model. Significance in the confusion matrices was assessed separately for each level of aggregation and data preparation method. Significance of difference in scanner TPR was assessed separately for sex and handedness data and sex, handedness, and age balanced data.

**Fig. 2.**
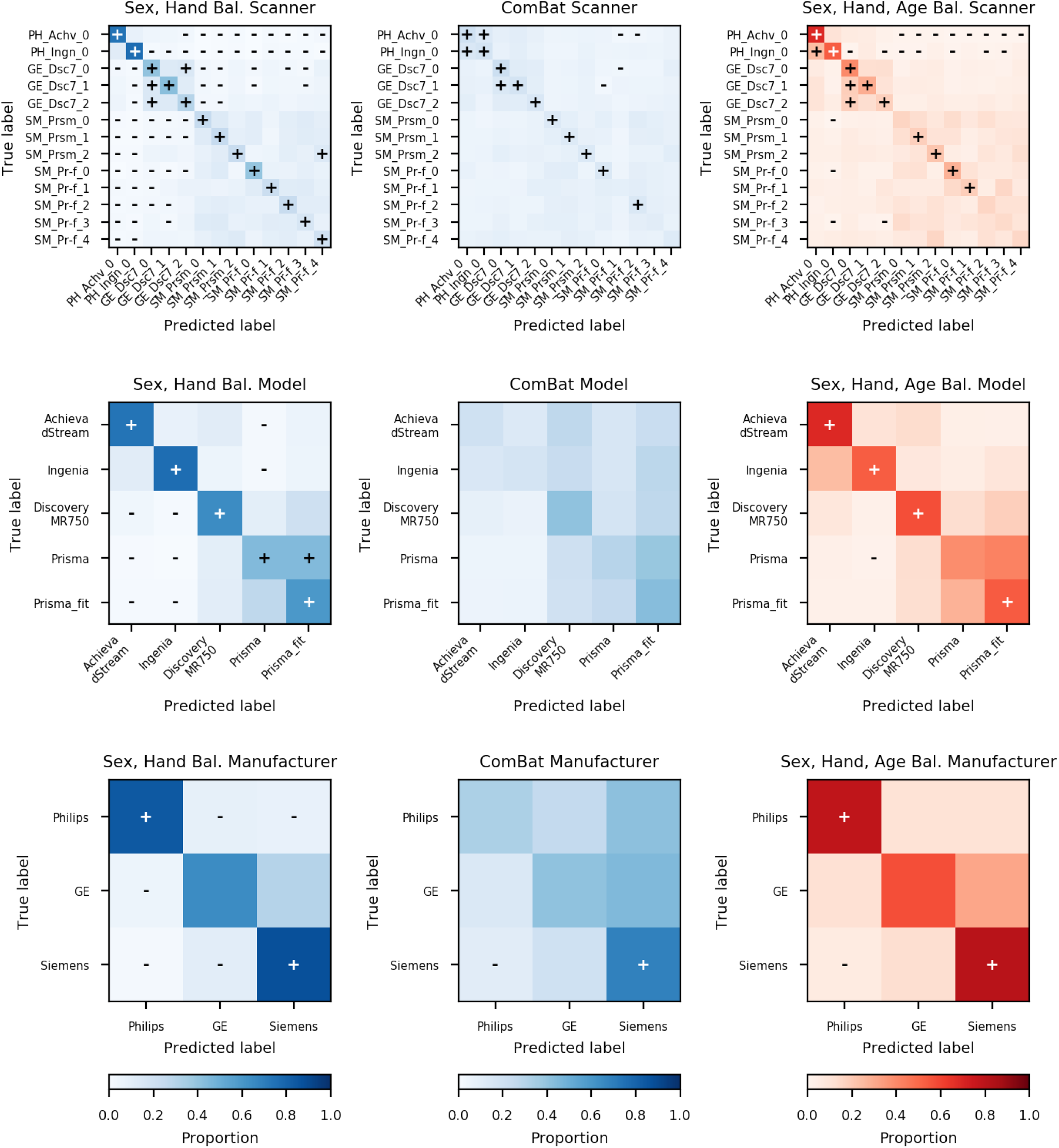
Confusion Matrices for Classification of Scanner with Resting State Data with Aggregation to Model and Manufacturer Levels. Intensity of the color in each cell represents the proportion of examples that were truly of that class the model predicted to be in that class. The diagonal represents correct classifications and the off-diagonal represents incorrect classifications. We ran 1,000 permutations balanced across gender, handedness and age in 3 month bins to determine chance classifier performance. Pluses and minuses indicate cells with significantly higher or lower values than expected by chance at a two-sided threshold of *p* < 0.005. Scanners are identified by abbreviation reflecting the manufacturer, model, and an arbitrary index number. The top row shows confusion matrices for classification of scanner from resting state data in test sets averaged across three-fold cross validation from 25 draws for the Sex, Handedness Balanced and ComBat corrected data and split-half cross validation from 25 draws for the Sex, Handedness, Age Balanced data. The second and third rows show these same results aggregated across scanners to the model and manufacturer levels. The Sex, Handedness Balanced Scanner matrix demonstrates that scanner was classifiable in the resting state data, particularly for the Philips and GE scanners and that Philips data was rarely classified as coming from a Siemens scanner. In the ComBat Scanner matrix we see that training ComBat separately on the training and test splits reduces classifiability of scanner, though the reduced performance was only below chance on three scanners. Scanner classification still performed above chance for nine scanners in the Sex, Handedness, Age Balanced data, demonstrating that classifiability of the resting state data was not driven by trivial differences in age distributions between scanners. Aggregating to the model level we see that scanner classification errors were more likely to occur between scanners using the same model of MRI scanner, though ComBat correction for scanner eliminated this tendency. Aggregating to the Manufacturer level demonstrates that scanner classification errors were also likely to occur between scanners using the same Manufacturer and that ComBat does not completely eliminate this tendency.

ABCD’s resting state connectivity consists of 412 connectivity metrics, as well as 6 measures of participant motion during the scan. We fit a set of ordinary least squares models to each of these metrics to determine the proportion of variance accounted for by scanner ID, model, and manufacturer, while controlling for variance accounted for by sex, handedness and age (Figure 3). 10 metrics have more than 10% variance accounted for by manufacturer, model, and scanner ID. ComBat correction for scanner eliminated significant manufacturer, model, and scanner variance for all metrics other than “Ventral-Ventralattn”, while preserving contributions to variance from sex, handedness, and age for all metrics. Motion parameters had small but significant contributions to variance from scanner and also varied with age as has been previously described (Power et al., 2012). Significance for the percent variance explained was calculated as a single tailed test with the null distribution constructed from the maximum percent variance explained across sex, handedness, and age for all 418 metrics for each permutation.

**Fig. 3.**
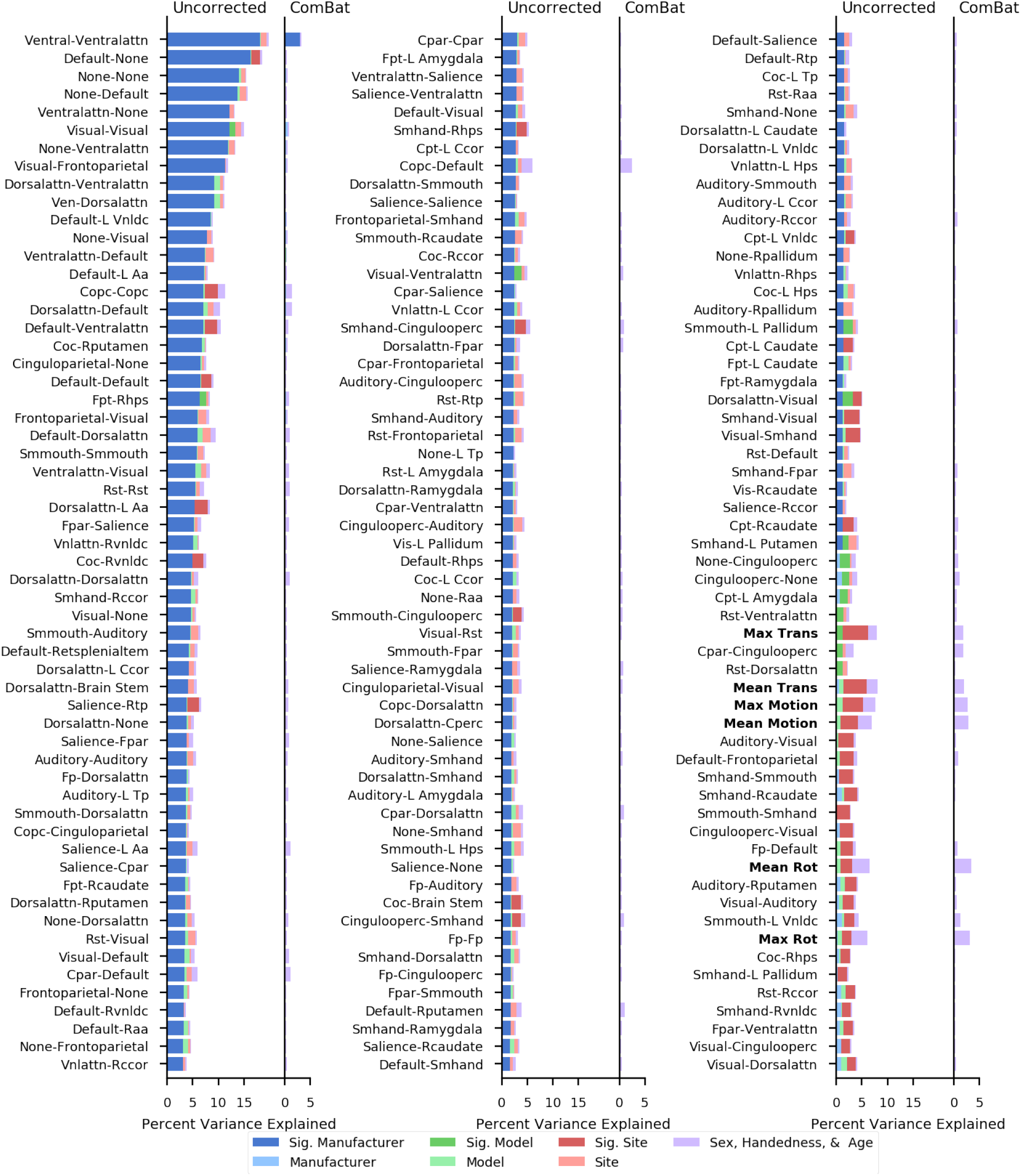
Sources of Variance Before and After Combat Correction for Resting State Functional Connectivity Between Networks and Subcortical ROIs. The 174 metrics for which one of manufacturer, model, or scanner was a significant source of variance are shown here. Significant sources of variance are indicated with more saturated colors. Significance was calculated individually for each of manufacturer, model, and scanner at *p* of 0.001 corrected for comparisons across connectivity metrics. Permutations preserved age, gender, and handedness distributions, so we cannot calculate the significance of these factors from these permutations. Motion parameters are identified with bold labels. ComBat correction for scanner eliminated significant contributions to variance from manufacturer, model, and scanner for all metrics except Ventral-Ventralattn, which refers to connectivity within the ventral attention network. All of the motion parameters had significant variance contributions from scanner as well as contributions from age. The 12 networks are: Auditory, Cingulo-Opercular, Cingulo-Parietal, Default, Dorsal Attention, Fronto-Parietal, Retrosplenial-Temporal, Salience, Somatomotor Hand, Somatomotor Mouth, Ventral Attention, and Visual. “None” refers to ROIs not assigned to a community.

### Task-Based fMRI

We classified collection scanner for ROI level within subject beta weights from each run of two runs for each of 26 contrasts or conditions in the ABCD task-based fMRI data (Figure 4). As with the resting state data we used a multinomial logistic regression with L2 normalization. The null distribution for the two tailed test of significance was determined from 1,000 sex, handedness and age balanced permutations and controlled for multiple comparisons across conditions/contrasts and ROIs as described above.

**Fig. 4.**
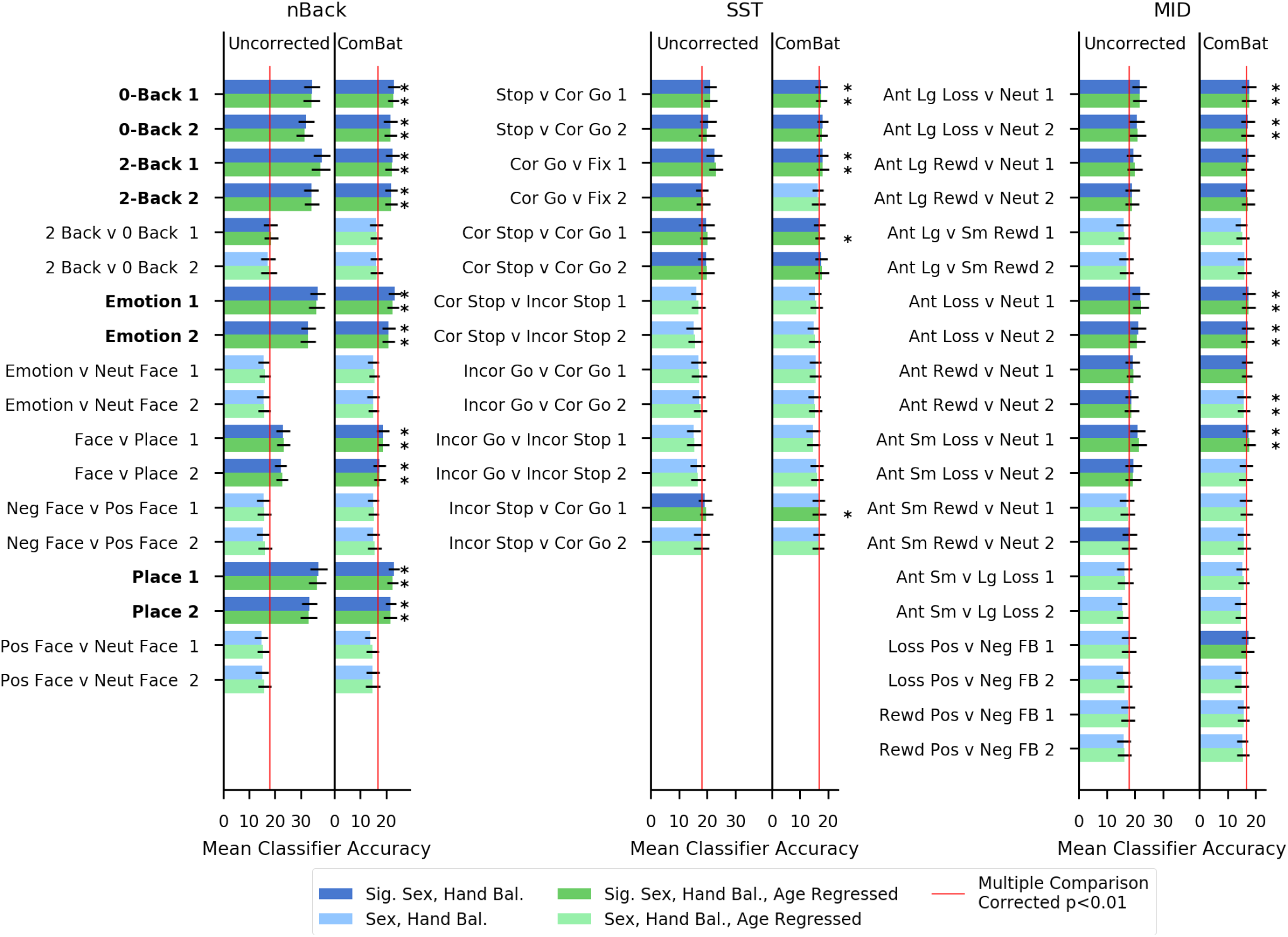
Classifier Performance Before and After ComBat for Task-Based fMRI Conditions and Contrasts. Bars show mean classifier performance across three-fold crossvalidation from 25 gender and handedness balanced draws for each contrast or condition from each task-based fMRI run. Errorbars indicate standard deviation across draws and cross-validation folds. Bold labels denotes conditions as opposed to contrasts. The number after each condition or contrast indicates run. We ran 1,000 permutations balanced across sex, handedness and age in 3 month bins to determine chance classifier performance. The red lines indicate the multiple comparison corrected *p* < 0.002 threshold for chance performance. For the uncorrected bars the threshold is 18.1% for ComBat it is 16.8%. Bars with performance greater than this cut off are indicated with more saturated colors. Classifier performance on data with no corrections for age are shown in blue. Green bars show classifier performance in which the effect of age learned on the training data for each split was regressed out of each ROI. The uncorrected columns show results without running ComBat to correct for scanner effects. The ComBat columns show classifier performance after correcting for scanner effects with ComBat based on parameters learned from the training split within each cross validation. It is notable that classifier performance is better for conditions than for contrasts, with significant contrasts performing just barely better than chance. Stars indicate that ComBat significantly reduced classifier performance with *p* < 0.002 threshold. SST: Stop Signal Task, MID: Monetary Incentive Delay.

At a threshold of *p* < 0.002, 9 contrasts spread across all 3 tasks were more classifiable than chance for both runs when regressing out the effect of age. The chance performance thresholds were 18.1% for uncorrected data and 16.8% for ComBat corrected data. The ABCD data release only provided beta weights on conditions for the nBack task. All eight runs from these four conditions were more classifiable than chance *p* < 0.002 and more classifiable than any of the contrasts with *p* < 0.002. ComBat reduced the classifier performance below chance for all of the contrasts and conditions. Considering the variance explained for each ROI from each task-based condition we see several ROIs have significant variance explained by scanner across all runs and conditions (Figure 5). There are also some ROIs with variance explained by manufacturer, but none across all conditions and runs. 19 ROIs have significant variance explained by scanner for multiple MID runs and contrasts (Figure 6). Two ROIs have significant variance explained by scanner model for two contrasts, “MID anticipated small loss versus neutral run one” and “MID anticipated loss versus neutral run one”. There are three ROIs with significant manufacturer variance for the “nBack Face versus Place” contrast and two ROIs with significant manufacturer variance for the “SST stop versus correct go” contrast.

**Fig. 5.**
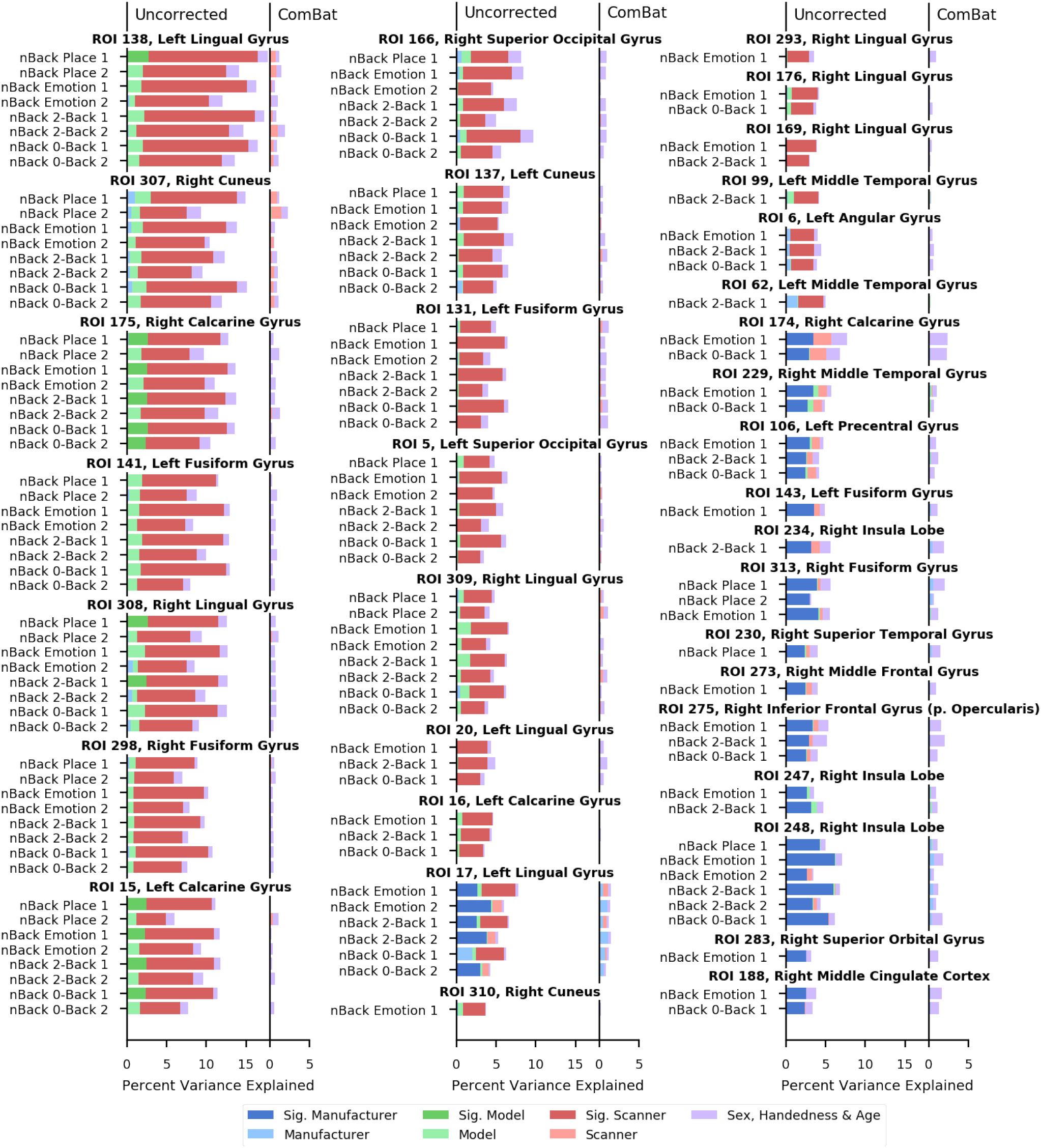
Sources of Variance in Different ROIs Before and After ComBat for Task-Based fMRI Conditions. Significant sources of variance are indicated with more saturated colors for manufacturer, model, and scanner. Significant percent variance explained is calculated individually for each of manufacturer, model, and scanner at a p-value threshold of 0.001 corrected for comparisons across ROIs. Permutations preserved age, gender, and handedness distributions, so we cannot calculate the significance of these factors from these permutations. Bars are grouped by ROI and labeled with the hemisphere and ROI number. SST: Stop Signal Task, MID: Monetary Incentive Delay.

**Fig. 6.**
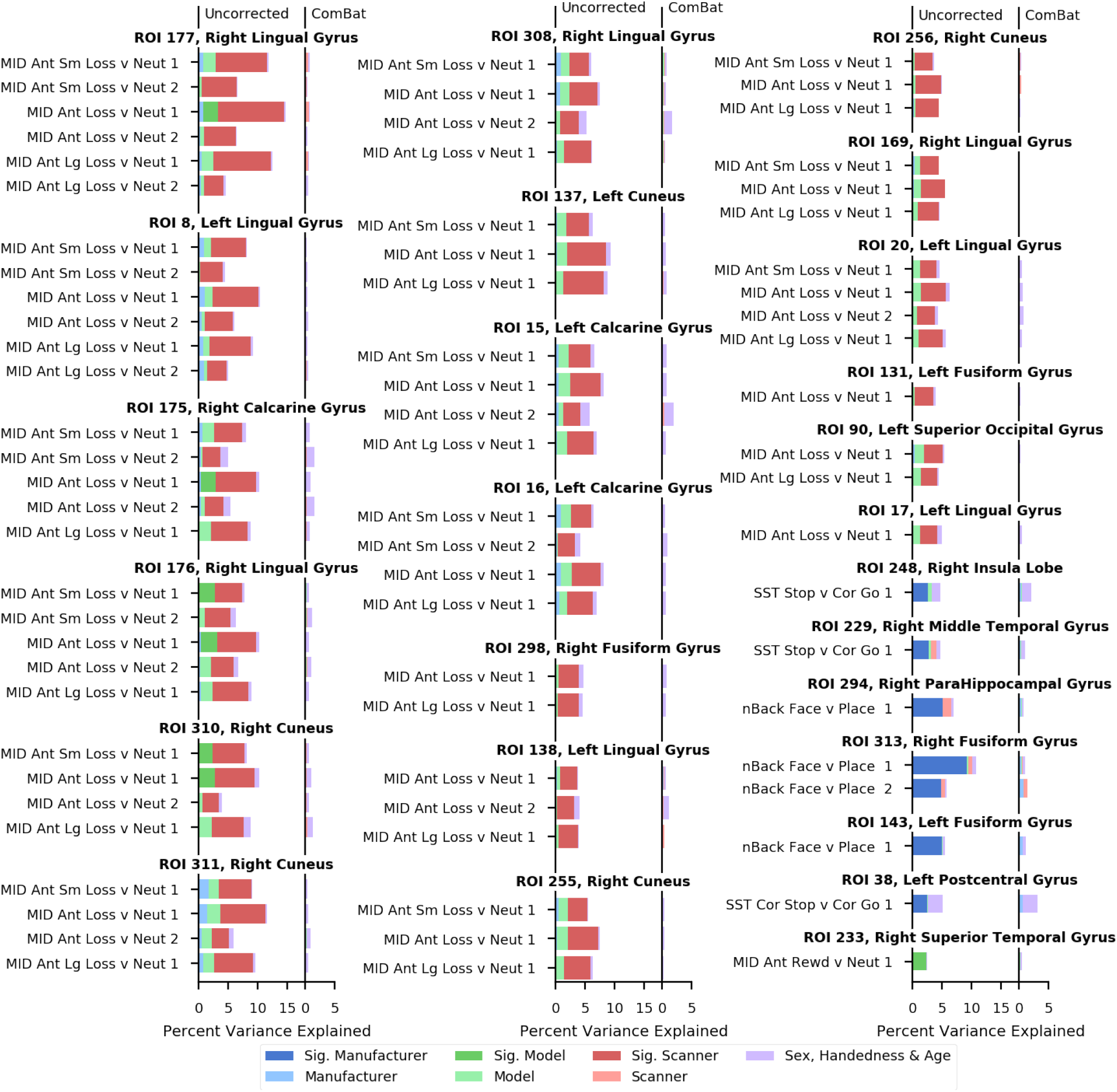
Sources of Variance in Different ROIs Before and After ComBat for Task-Based fMRI Contrasts. Significant sources of variance are indicated with more saturated colors for manufacturer, model, and scanner. Significant percent variance explained is calculated individually for each of manufacturer, model, and scanner at a p-value threshold of 0.001 corrected for comparisons across ROIs. Permutations preserved age, gender, and handedness distributions, so we cannot calculate the significance of these factors from these permutations. Bars are grouped by ROI and labeled with the hemisphere and ROI number.SST: Stop Signal Task, MID: Monetary Incentive Delay.

We also looked for ROIs with consistent coefficients across the nBack conditions as a way to assess the spatial distribution of scanner differences (Figure 7). We found coefficients for three ROIs across one GE scanner and two Siemens scanners that were consistently different from zero across conditions and runs at a *p* < 0.002. These coefficients were exclusively for occipital ROIs.

**Fig. 7.**
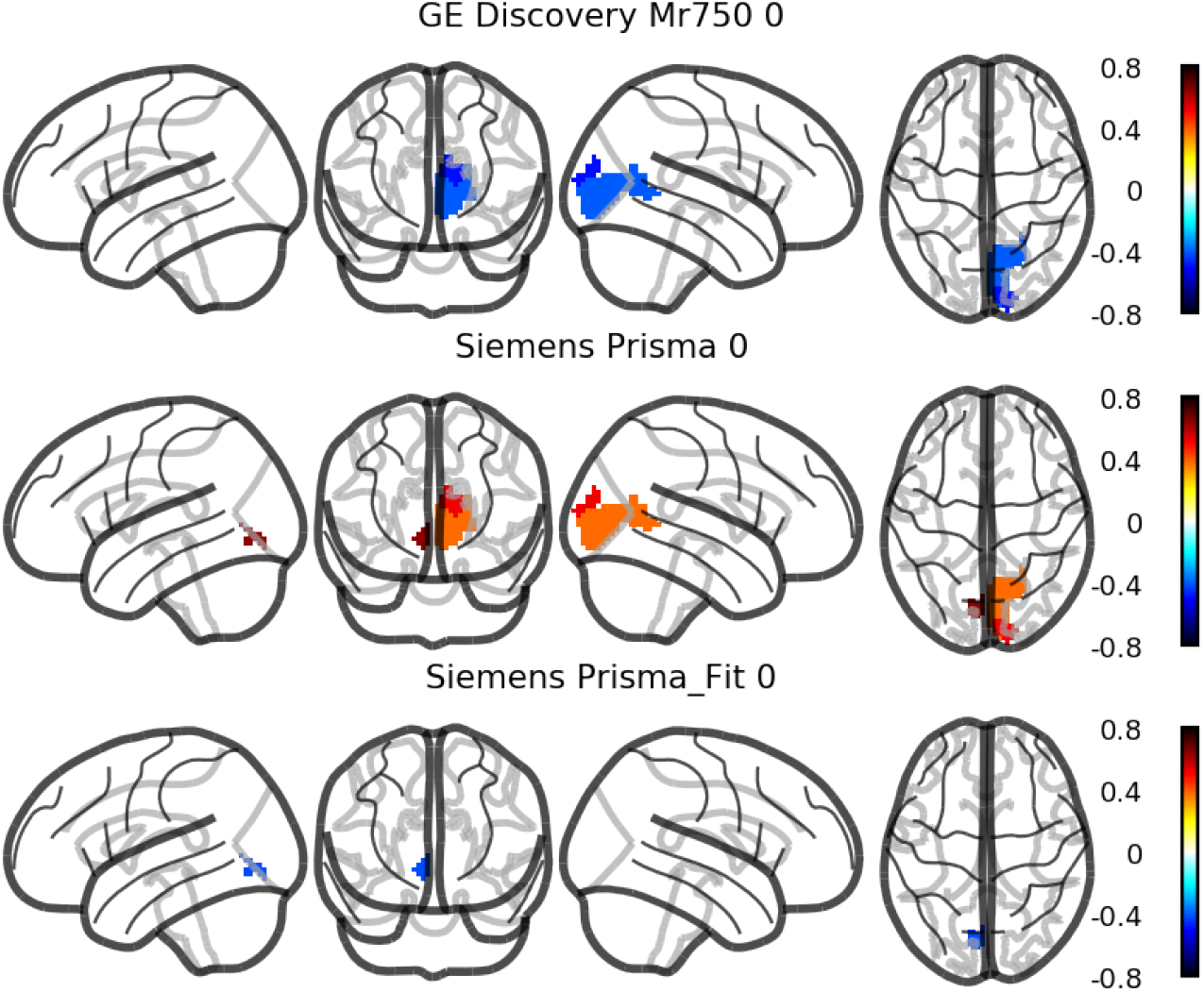
Spatial Distribution of Coefficients Contributing to Scanner Classifiability during nBack Conditions. Colors reflect the average value of multinomial logistic regression coefficients for models classifying scanner across two runs each of the four nBack conditions. Since we fit a multinomial logistic regression, we have a coefficient for every ROI at scanner. The ROI level coefficients, plotted in MNI space, provide insight into the spatial distribution of differences contributing to scanner classifiability. The ROIs driving scanner differences across conditions are in the occipital lobe. Only coefficients that are consistent across runs and conditions are shown. Consistency was assessed as significant difference from zero at a threshold of *p* < 0.002 across runs and conditions, with the null distribution correcting from multiple comparison coming from 1,000 age, sex, and handedness balanced permutations. Four of the seven scanners did not have any consistent coefficient and are therefore not shown. Left is plotted on the left.

## Discussion

In this manuscript we have quantified variance associated with scanner manufacturer, model, and serial number in the ABCD Study’s rs-fMRI and task-based data. Four scanners (two Philips and two GE) have the greatest classifiability in the resting state data. Classification mistakes are more likely between scanners from the same manufacturer and scanners with the same models. Ten resting state metrics have more than 10% variance explained by manufacturer, model, and scanner ID. Motion parameters also differ by scanner but not by manufacturer, indicating that there may be site differences in subject preparation.

We have also shown that ComBat (Johnson et al., 2007; Fortin et al., 2018), a recently proposed method for removing unwanted source of variance using Empirical Bayes, effectively removes the majority of scanner related variance. This is demonstrated in both our classifier experiment and when examining the percent variance explained for each metric in both the rs-fMRI and task-based data. Even though we only modeled and removed scanner with ComBat, it also effectively removed variance due to scanner model and manufacturer for all but one variable in the rs-fMRI data. ComBat as currently implemented cannot control for more than one variable at a time. It may be possible to do so through sequential application of ComBat, but we have not evaluated this application.

While we were able to classify scanner above chance for both conditions and contrasts in the Task-based fMRI, the conditions are much more classifiable than the contrasts. The subtraction involved in computing the contrasts may be sufficient to largely remove variance due to scanner, which is shared between the two conditions. Since the task-based data is provided at the ROI level, the ROIs that are consistently contributing to classifiability can give us some insight into the spatial distribution of scanner differences. We find consistent contributions to scanner differences across condition within the occipital lobe. Differences in the occipital lobe may be due to difference in stimulus hardware (luminance and contrast of display, etc.), or simply a result of the occipital lobes response to visual stimuli across all of the nBack conditions. Investigating conditions from additional tasks may provide more insight into the spatial distribution of scanner differences. These may be coming from differences in EPI distortion, dropout, or registration.

Previous studies examining cross-scanner effects have explored the effect of cross-scanner variance on specific hypotheses being investigated (Focke et al., 2011; Auzias et al., 2016). The ABCD study is primarily designed to investigate the interaction of drug use with neurodevelopment, but the data released thus far is only from the first time point of the longitudinal study. Potential participants who had already used these substances were excluded from the study, so the data released thus far does not contain any information about these contrasts yet. Beyond that, given that the ABCD study is making a large, longitudinal study of neurodevelopment publicly available, it is likely that many additional hypotheses will be tested with this dataset. Since we could not evaluate cross scanner effects relative to ABCD’s primary hypothesis and do not know what other hypotheses may be tested with this data, the variance explained by sex, handedness, and age in months must serve as stand-ins for determining the impact of cross-scanner effects. If you are considering an analysis with the ABCD data and are trying to decide the extent to which your hypothesis may be affected by cross-scanner effects, imagine the magnitude of the effect you expect to observe relative to the effect of sex, handedness and age. If you expect your effect of interest to have about the same magnitude as the effect of sex, handedness and age, then crossscanner differences are likely to be at least the same order of magnitude as your effect for many of the rs-fMRI metrics, as well some of the task-based metrics. ComBat does provide an effective means of reducing scanner related variance and we recommend that other groups analyzing imaging data from cross-scanner data sets consider including it in their analyses.

## Conclusions

Scanner effects exist in metrics for rs-fMRI and task-based conditions released by the ABCD study, but task-based contrasts show minimal scanner effects. This implies that other metrics that also have a within-dataset subtraction may also be somewhat insulated from cross-scanner effects, but we have only analyzed ABCD’s functional data in this paper. ComBat performs well at removing scanner effects and should be considered for inclusion in cross-scanner analysis pipelines.

## ACKNOWLEDGEMENTS

**ABCD Data.** Data used in the preparation of this article were obtained from the Adolescent Brain Cognitive Development (ABCD) Study (https://abcdstudy.org), held in the NIMH Data Archive (NDA). This is a multi-site, longitudinal study designed to recruit more than 10,000 children age 9-10 and follow them over 10 years into early adulthood. The ABCD Study is supported by the National Institutes of Health and additional federal partners under award numbers U01DA041022, U01DA041028, U01DA041048, U01DA041089, U01DA041106, U01DA041117, U01DA041120, U01DA041134, U01DA041148, U01DA041156, U01DA041174, U24DA041123, and U24DA041147. A full list of supporters is available at https://abcdstudy.org/nih-collaborators. A listing of participating sites and a complete listing of the study investigators can be found at https://abcdstudy.org/principal-investigators.html. ABCD consortium investigators designed and implemented the study and/or provided data but did not necessarily participate in analysis or writing of this report. This manuscript reflects the views of the authors and may not reflect the opinions or views of the NIH or ABCD consortium investigators.

The ABCD data repository grows and changes over time. The ABCD data used in this report came from doi://10.15154/1412097. DOIs can be found at http://dx.doi.org/10.15154/1412097.

**HPC.** This work utilized the computational resources of the NIH HPC Biowulf cluster. (http://hpc.nih.gov)

